# Predicting division planes of three-dimensional cells by soap-film minimization

**DOI:** 10.1101/199885

**Authors:** Pablo Martinez, Lindy A. Allsman, Kenneth A. Brakke, Christopher Hoyt, Jordan Hayes, Hong Liang, Wesley Neher, Yue Rui, Allyson M. Roberts, Amir Moradifam, Bob Goldstein, Charles T. Anderson, Carolyn G. Rasmussen

**Affiliations:** Center for Plant Cell Biology, Department of Botany and Plant Sciences, University of California, Riverside; Biochemistry and Molecular Biology Graduate Program, University of California, Riverside; Department of Mathematics, Susquehanna University, Selinsgrove, PA; Center for Plant Cell Biology NSF-REU, Harvey Mudd College; Institute of Integrative Genome Biology, University of California, Riverside; Department of Biology, The Pennsylvania State University; Curriculum in Genetics and Molecular Biology, University of North Carolina at Chapel Hill; Department of Biology, University of North Carolina at Chapel Hill; Department of Mathematics, University of California, Riverside

## Abstract

One key aspect of cell division in multicellular organisms is the orientation of the division plane. Proper division plane establishment contributes to normal organization of the plant body. To determine the importance of cell geometry in division plane orientation, we designed a threedimensional probabilistic mathematical modeling approach to directly test the century-old hypothesis that cell divisions mimic “soap-film minima” or that daughter cells have equal volume and the resulting division plane is a local surface area minimum. Predicted division planes were compared to a plant microtubule array that marks the division site, the preprophase band (PPB). PPB location typically matched one of the predicted divisions. Predicted divisions offset from the PPB occurred when a neighboring cell wall or PPB was observed directly adjacent to the predicted division site, to avoid creating a potentially structurally unfavorable four-way junction. By comparing divisions of differently shaped plant and animal cells to divisions simulated in silico, we demonstrate the generality of this model to accurately predict in vivo division. This powerful model can be used to separate the contribution of geometry from mechanical stresses or developmental regulation in predicting division plane orientation.

## Introduction

Cell division planes are dictated by geometric, mechanical and polarity cues in plants, animals, bacteria, and fungi (Minc and Piel, 2012). A challenging problem in understanding division plane orientation lies in separating the effects of cell polarity or mechanical cues from the effects of cell shape-mediated cues. In plant and animal cells, the absence of external polarity or mechanical cues often leads to a division plane which bisects the long axis of the cell (Besson and Dumais, 2014; Minc and Piel, 2012; Errera, 1888). In zebrafish embryos, the placement of future divisions can be predicted by cell shapes (Xiong et al., 2014). In the late 1800s, biologists identified basic patterns of plant cell division. The plane of division is typically perpendicular to the primary growth axis of the tissue (Hofmeister, 1863). The new cell wall often forms at a 90 degree angle to the mother cell wall (Sachs, 1878). Plant cell divisions appear to mimic soap films, often dividing along the smallest local plane to minimize the surface area of the division (Errera 1888, Besson and Dumais 2014). Later oversimplification from multiple planes to a single global minimum division plane significantly limited the ability to account for the observed variability in division plane orientation, leading biologists to ignore this problem for decades (Besson and Dumais, 2014).

Recently, researchers have used computational or mathematical approaches to understand division plane orientation in plant cells in 2 dimensions (2D) (Dupuy et al., 2010; Besson and Dumais, 2011; Sahlin and Jönsson, 2010). In several studies, empirically derived factors were added to account for the stochasticity of the observed division orientations (Dupuy et al., 2010; Besson and Dumais, 2011). The length difference between two predicted divisions, with addition of an empirically defined stochasticity factor, was sufficient to describe the relative proportions of population level divisions in cells from several plant species (Besson and Dumais, 2011). Other 2D approaches modeled different division plane preferences without using stochasticity in the *A. thaliana* shoot apical meristem. The shortest path through the center of mass of the cell best fit the observations, although it incompletely captured in vivo size variability (Sahlin and Jönsson, 2010). A fitness function that combined length minima for new cell walls with daughter cells of equal areas accurately predicted division planes and functioned similarly to “modern” Errera predictions (Shapiro et al., 2015).

An interest in 3D modeling of cell division led to division plane analysis in the *A. thaliana* embryo (Yoshida et al., 2014). The center of mass for each cell was used as a point to sample 2000 different planes to identify the lowest flat surface area. Some embryonic cells did not divide according to the shortest plane, but instead divided asymmetrically to produce unequal daughter cell volumes. Asymmetric divisions in the embryo were driven by response to auxin and associated with alterations in both gene expression and differentiation. Mutants that do not respond to auxin lost division asymmetry in these cells (Yoshida et al., 2014). While this approach did not minimize surface areas locally or provide a probabilistic prediction of division plane orientation it was successfully used to predict a potential global minimum in 3D.

Computational approaches have begun modeling the dynamics of interphase microtubule arrays using 3D shapes with a potential long-term application of predicting division plane orientation. Modeling microtubule properties such as directionality, interactions via crosslinking proteins or interactions with the cell wall were sufficient to promote in silico localization of microtubules to the cortex of a 3D simulated plant cell (Mirabet et al., 2018). The calculated microtubule array depended on cell shape cues but could also be modulated by external forces (Mirabet et al., 2018). Changing either microtubule dynamics or specific face or edge properties generated cortical microtubule arrays in realistically shaped cells (Chakrabortty et al., 2018a). Understanding how the cortical microtubule array may be oriented by cell shape and other parameters might help predict the orientation of the future division plane. This model of cortical microtubule arrays in Arabidopsis embryo cells was applied to generate cortical microtubule arrays aligned with division planes, but they were not compared to in vivo divisions (Chakrabortty et al., 2018b).

Accurate division plane prediction provides a mechanism to weigh relative contributions of multiple, potentially developmentally or mechanically regulated, drivers of plant growth. Here, we used a 3D mathematical modeling approach to generate multiple division-plane predictions for any cell. This 3D model explicitly tests the long-held hypothesis that plant cell divisions mimic soap-film minimum surfaces (Errera, 1888). This model depends only on the shape of the cell: division predictions are performed by initiating soap-film-like, area-minimizing, descending gradients from starting planes designed to fully and evenly sample the volume of the cell. This geometry-based model identifies cases when plant cells divide according to their geometry, and highlights when they do not. The location of PPBs, microtubule and microfilament structures that accurately predict the future division plane in typical land plant cells (Martinez et al., 2017; Pickett-Heaps and Northcote, 1966; Camilleri et al., 2002; Van Damme et al., 2007; Smertenko et al., 2017), most often closely match one class of the predicted division planes. Discrepancies in PPB location compared to predicted divisions was sometimes due to PPB shifting in response to an adjacent cell wall. Finally, we demonstrate that the model provides accurate predictions for the location of the future division of diverse symmetrically dividing cells including maize epidermal cells and developing ligule cells, *Arabidopsis thaliana* guard cells, and *C. elegans* embryonic cells.

## Results

We established a geometry-based model to generate local minimum (soap-film) predicted divisions for any cell shape using Surface Evolver (Brakke, 1992). First, confocal micrographs of maize cells with PPBs were taken with 0.2 or 0.4 micron-interval Z-stacks (Figure 1A-C shows a single Z-plane). The images were semi-automatically thresholded and their 3-dimensional (3D) surfaces were extracted using FIJI (Figure 1D, Materials and Methods). The surfaces were imported into Surface Evolver. The surfaces were smoothed using 30th degree spherical harmonics, a method which approximates the cell outline as a function of polar coordinate angles, analogous to Fourier transformation approximations of 2D shapes (Figure 1F, 1J-K, Supplemental Figure 1, Materials and Methods) (<u>Givoli, 2004</u>) and commonly used in 3D rendering (<u>Shen et al., 2009</u>). Next, mathematically predicted divisions were generated for each of the cells using a set of 241 starting planes which evenly sampled the entire cell volume (Materials and Methods). These starting planes were used to initiate a process known as gradient descent. Gradient descent iteratively minimized the initial surface area until the lowest local surface area (soap-film minimum) was reached to divide the mother cell into two daughter cells with equal volume (Figure 1G).

**FIG. 1.**
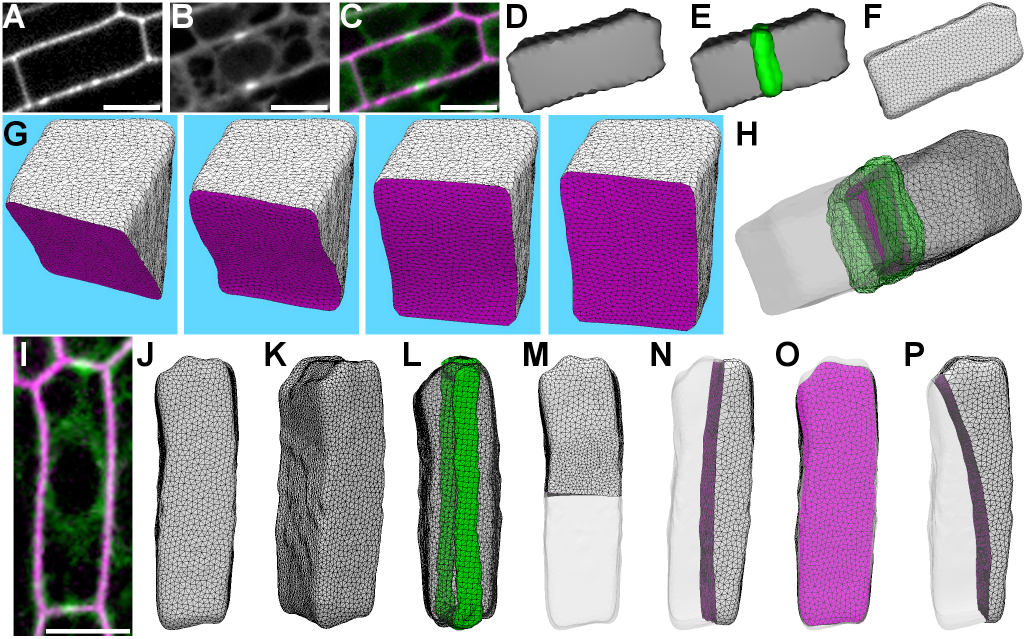
Division plane predictions and their comparison to in vivo divisions. (A) Maize adaxial epidermal cell wall stained with propidium iodide. (B) Same cell expressing YFP-TUBULIN to identify the PPB. (C) Merged image of cell wall (magenta) and PPB (green). (D) Reconstruction of the 3D surface. (E) The 3D surface overlayed with the 3D PPB. (F) Surface generated in Surface Evolver after 30th degree spherical harmonics. (G) An example of gradient descent identifying the soap-film local minimum from an initial starting position (left panel) to an anticlinal transverse division plane. (H) The PPB (green) colocalized with one of the final predicted divisions (magenta). (I) Maize leaf epidermal cell wall stained with propidium iodide (magenta) and expressing YFP-TUBULIN (green) with a longitudinally oriented PPB. (J-L) 3D cell surface of example shown in (I). (J) Cell oriented as shown in micrograph from (I). (K) Cell oriented to show surface along the z-axis. (L) Cell surface with PPB (green). (M-P) Examples of each predicted division class generated by identification of local minimum surfaces. (M) transverse division plane, the global minimum division surface. (N) longitudinal division (O) periclinal division. (P) “other” division. Scale bars are 10μm.

The local minima identified from 241 independent soap-film minimizations tended to coalesce on discrete locations within the cell. The final predicted surface areas of division planes were closely matched in size and location. Soap-film minimization of rectangular prism-shaped epidermal cells from two different maize leaf developmental stages generated between 1 and 4 local minima. Based on the location of the predicted division and its location within the cell relative to the tissue, these local minima were classified as transverse (Figure 1M), longitudinal (Figure 1N), periclinal (Figure 1O) and other (Figure 1P). The most common predicted division was a transverse, anticlinal division which was predicted for all cells (n = 181), followed by periclinal divisions (98%, 178/181) and longitudinal anticlinal divisions (63%, 114/181). The majority of maize epidermal cells (78%, n = 141/181) displayed a PPB which closely spatially matched one of the predicted divisions. A small number of cells did not have any predicted division which matched the orientation of the PPB even broadly (2%, n= 4/181), while 20% (n= 36/181) had small spatial discrepancies discussed in detail in two paragraphs.

For the majority of maize epidermal cells (78%, n = 141/181) the PPB closely spatially matched one of the predicted divisions. YFP-TUBULIN signal was used to reconstruct the the PPB in 3D (Figure 1E, 1L). Next, the PPB was aligned with the predicted surface closest to the PPB (Figure 1H). To quantitatively assess PPB and predicted division overlap, we measured the distance in μm^2^ between the midplane of the PPB and the cortical region of the predicted division. As the distance reaches 0, the predicted division more closely aligns with the PPB midplane. The average distance or offset between the PPB and the closest predicted division was 0.492 +/− 1.156 μm^2^; mean +/− standard deviation (S.D.), n = 141/177 (the 4 cells where the PPB and prediction had no overlap were not included; Figure 2J). These data indicate that the predicted division typically localized within the boundary of the PPB.

**FIG. 2.**
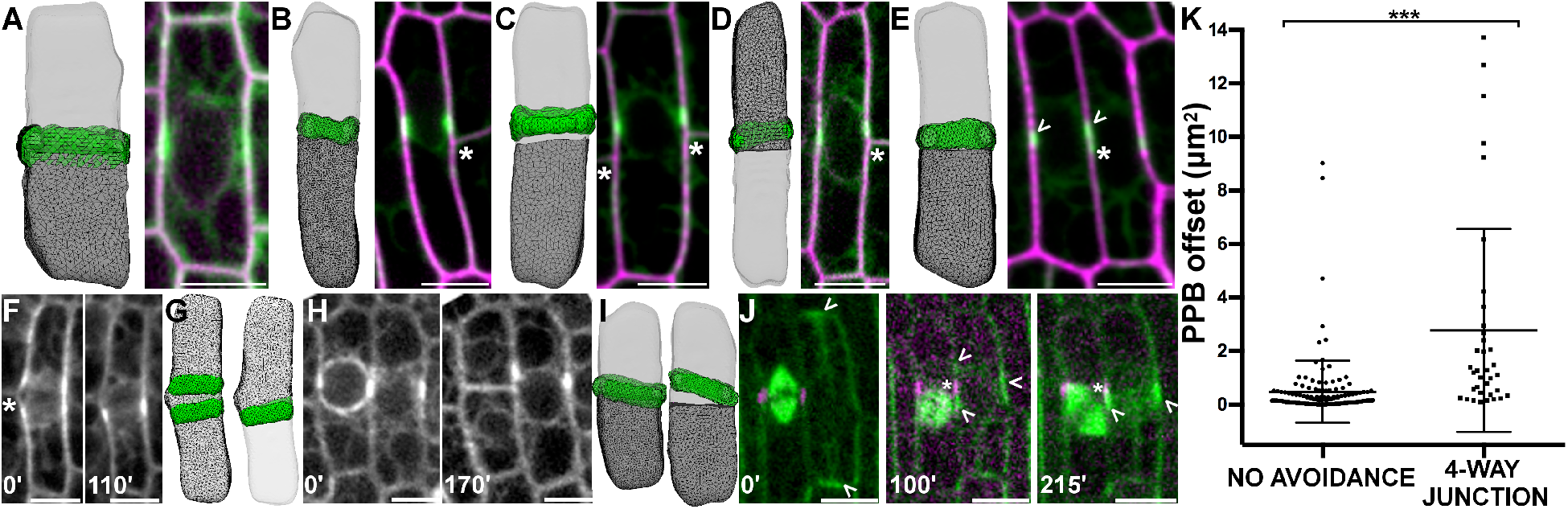
Comparison between predicted divisions and location of the PPB: Identification of 4-way junction avoidance. (A-E) Best-fit predicted divisions overlaid with in vivo PPB location compared to in vivo maize epidermal cells expressing YFP-TUBULIN (green) and either expressing membrane marker PIP2-CFP (A,C,D; magenta) or stained with propidium iodide to outline the cell wall (B,E; magenta). Neighboring cell wall or PPB indicated with an asterisk. (A) Cell with low PPB offset, 0.04 μm^2^. (B) Cell where the predicted division and the PPB are slightly offset at 0.54 μm^2^. (C) Example of cell with two neighboring cell walls and a high offset between the PPB and the predicted division (12.69 μm^2^). (D) Cell with neighboring cell wall with high PPB offset in one direction (13.72 μm^2^). (E) Cell which has an offset PPB caused by a neighboring PPB (0.10 μm^2^). PPB marked by arrowheads. (F-H) Timelapse of cells expressing YFP-TUBULIN where neighboring cells potentially influence PPB location. (F) Cell with a split PPB (left) that collapsed into one PPB before entering mitosis (time in minutes on the lower left). (G) Reconstruction of cell surface with both possible PPBs next to best fit predicted division to the PPB which was chosen (offset = 0.13 μm^2^). (H) Cell (left) with a PPB that initiated mitosis next to a neighboring cell still in pre-prophase with an offset PPB. (I) Reconstruction of both cell surfaces with the best fit predicted division overlaid onto each respective PPB, offset for cell on left is 0.13 μm^2^ while the offset for the cell on the right is 5.59 μm^2^. (J) Timelapse of cell expressing CFP-TUBULIN (green) and TAN1-YFP, which labels the division site (magenta). The cell on the right had a longitudinal PPB that switched to a split transverse PPB without signal directly adjacent to the division site of the neighboring cell. (K) PPB offset (distance from the midplane of the PPB to the closest predicted division surface) was 0.492 +/− 1.156 μm^2^ S.D. (S.D. n = 136/177) for cells with PPBs without neighbor cell walls or PPBs near the predicted in silico division plane. Cells with predicted divisions that would form a 4-way-junction (due to an adjacent neighbor wall or PPB) had significantly higher offset between the predicted division and the PPB location (average 2.778 +/− 3.793 μm^2^ S.D., n = 36/177, Mann-Whitney *P* = < 0.0001). Scale bars are 10μm.

In a subset of the cells (~20% n = 36/181), small, but noticeable deviations between PPB location and predicted divisions were observed, likely due to neighboring cell influence.

The PPB was visibly and significantly offset from the in silico predicted division plane to avoid creating a four-way junction with an already established adjacent cell wall next to the predicted division plane (offset = 2.778 +/− 3.793 μm^2^; mean +/− S.D. (Figure 2K). PPBs adjacent to sites of neighboring cell influence, either an adjacent cell wall (Figure 2B-D, n = 34) or an adjacent PPB (Figure 2E, n = 2), were shifted from the mathematically optimal division plane on average five times further than PPBs without obvious neighboring cell influence. Avoidance of four-way junctions is a recognized feature of plant cells thought to increase structural stability (Flanders et al., 1990). In vacuolate cells, dense rings of cytoplasm, called phragmosomes, accumulate at the division site. Neither phragmosomes in vacuolate plant cells (<u>Sinnott and Bloch, 1941</u>) nor PPBs (<u>Gunning et al., 1978</u>) in adjacent cells overlap, but, to the best of our knowledge, this is the first demonstration that cells with neighboring cell walls shift the PPB away from a soap-film minimum predicted division plane. It is not currently known how four-way-junction avoidance occurs.

We used time-lapse imaging to analyze PPB movement and avoidance in live cells, revealing the dynamic nature of PPBs and their interaction with neighboring cells. Dynamic PPB events such as avoidance of adjacent cell PPBs or cells were rare (n=6, <1% of forming PPBs observed during time-lapse imaging) limiting the sample size. Time-lapse imaging revealed a PPB split into two microtubule accumulations across an adjacent cell wall (Figure 2F, time 0) that later coalesced into one location (Figure 2F-G, time 110’). Time-lapse images showed two non-overlapping but adjacent PPBs. One PPB closely matched the soap-film minimum while the other had high offset that was maintained as one neighboring cell completed mitosis (Figure 2H-I). Finally, we observed a PPB shifting from longitudinal (Figure 2J, time 0’) to transverse, while splitting into two segments to avoid an adjacent cell division site indicated by TAN1-YFP localization (Figure 2J, time 100’), then finally coalescing on one position (Figure 2J, time 215’).

These data demonstrated that PPB shifting occurred in vivo when some interaction with an adjacent cell promoted PPB offset.

Longitudinal and transverse symmetric divisions were observed in vivo during two distinct maize leaf developmental stages (described below) via PPB location. Cells with longitudinal PPBs (n = 36) had significantly higher proportions of predicted longitudinal divisions after soap-film minimization (predicted longitudinal division frequency = 5.05% +/− 3.61; mean +/− S.D.) compared to cells with transverse PPBs (n = 145, predicted longitudinal division frequency 1.85% +/− 2.60; mean +/− S.D., Mann-Whitney *P* = <0.0001, Figure 3A). To determine whether there was a characteristic feature of longitudinally or transversely dividing cell shapes, we used a metric independent of cell size called the eigenvalue ratio (described in materials and methods) to evaluate shape features. As the eigenvalue ratio approaches one, the cell appears less prolate, or more cube shaped. Longitudinally dividing cells had significantly lower eigenvalue ratios than transversely dividing cells (Figure 3B). Cells with lower eigenvalue ratios were more cube-shaped, while cells with higher ratios tended to be long and thin, more prolate (Figure 3C). Calculated eigenvalue ratios for each cell were plotted as a function of percent transverse predicted divisions: as cells become more cube-shaped, less transverse division predictions and more longitudinal and periclinal divisions were predicted (Figure 3D). Although a direct comparison cannot be made, this result is similar to elegant cell-shape based division predictions made for 2D cells in which the probability of the predicted division was more equally shared if the division lengths were similar in size (Besson and Dumais, 2011). This result suggests that cells that will divide longitudinally tend to be less prolate or more cube-shaped.

**FIG. 3.**
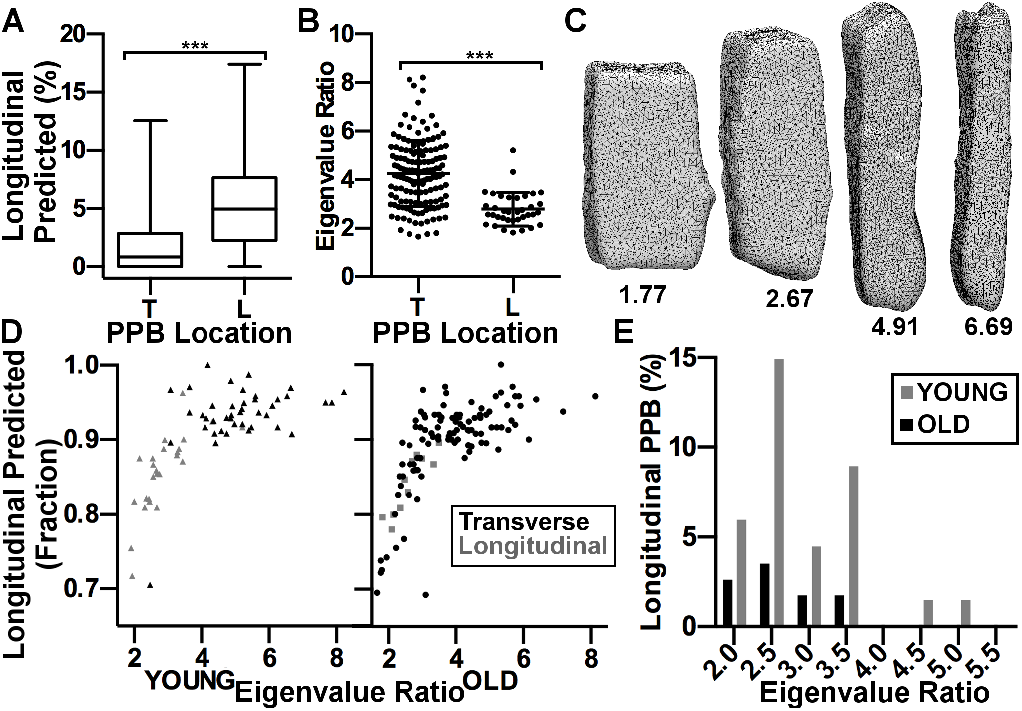
Cells with longitudinal PPBs were more cube-shaped and had more longitudinal predictions in silico than cells with transverse PPBs. (A) Box and whisker plot of the percent longitudinal divisions predicted between transverse or longitudinal PPBs. Cells with longitudinal PPBs (n = 36) predicted significantly (mean = 5.05% +/− 3.61 S.D.) more longitudinal divisions than those with transverse PPBs (n = 145, mean = 1.85% +/− 2.60 S.D., Mann-Whitney, *P* = <0.0001). (B) Dot plot of eigenvalue ratios of cells with a transverse (n = 145, mean = 4.26 +/− 1.36 S.D.) or longitudinal PPB (n = 36, mean = 2.80+/−0.70 S.D.). Cells with transverse PPBs had higher eigenvalue ratios (Mann-Whitney, *P* = <0.0001). (C) Cells with varying eigenvalue ratios. Larger eigenvalue ratios reflected longer and thinner cells (cells not displayed to scale). (D) Eigenvalue ratio versus fraction transverse division predicted. Cells with higher eigenvalue ratios ended up having higher proportions of transverse divisions predicted, a relationship with significant correlation (*P* = <0.0001 for both leaves; young leaf Pearson R squared = 0.4627; Old leaf Pearson R squared = .4920). (E) Histogram of cells which display longitudinal PPB binned by eigenvalue ratio for both maize leaf datasets. The young leaf sample had higher proportions of cells with longitudinal PPBs across all eigenvalue ratios.

Two different developmental stages of the maize leaf were analyzed. As the maize leaf develops, the proportions of symmetric division classes change. Developmentally younger leaf epidermal cells have higher proportions of longitudinal divisions that decrease in frequency as the leaf grows (Sylvester, 2000). Adaxial epidermal cells from a single developmentally younger leaf (leaf L14, leaf height 15.58 mm from a 28 day-old maize plant) and a single developmentally older leaf (L9 or L10, ligule height ~2 mm from a 28 day-old maize plant) were analyzed. Cells with transverse anticlinal divisions were the most common in both maize epidermal cell developmental stages. While the younger leaf had 37% longitudinal and 63% transverse PPBs (n = 67), the developmentally older leaf had ~10% longitudinal and ~90% transverse PPBs (n = 114, Table 1). In both samples, cells with longitudinal PPBs had more predicted longitudinal divisions in silico (previous paragraph and Figure 3B). However, in vivo longitudinal divisions occurred significantly more often than predicted by Surface Evolver soap-film minimization (2.6% total longitudinal predictions in young leaf cells versus 37% in vivo and 2.3% longitudinal predictions in old leaf cells versus 10%; Table 1), indicating that soap-film minimization underpredicts longitudinal divisions in vivo. The young leaf cells had proportionally more longitudinal PPBs across all eigenvalue ratios potentially suggesting developmentally regulated shifts in the probabilities of division plane positioning in response to shape (Figure 3E). Our data indicate that in vivo population level division plane orientations of cells with the same shape do not occur with the same frequencies in different developmental contexts nor can they be fully predicted using a geometry-based soap-film model. This suggests that developmental or mechanical forces also play a role orienting the final division plane (Figure 3D).

**TABLE 1:** In-vivo division class percents and in-silico division predictions

| TISSUE | n<br>(cells) | In-vivo (%) |  |  | In-silico (mean +/- SD) |  |  |  |
| --- | --- | --- | --- | --- | --- | --- | --- | --- |
|  |  | Transverse | Longitudinal | Periclinal | Transverse | Longitudinal | Periclinal | Other |
| Old Leaf | 114 | 90.3 | 9.7 | 0 | 89.1 +/- 6.1 | 2.6 +/-3.0 | 8.0+/-3.9 | 0.3+/- 1.9 |
| Young Leaf | 67 | 62.7 | 37.3 | 0 | 90.5+/-6.0 | 2.3+/-3.3 | 6.8+/-3.4 | 0.5+/-2.5 |
| Ligule Region* | 34 | N/A | N/A | 100 | 17.5+/- 17.4 | 11.9+/-17.5 | 68.8+/- 22.9 | 1.8+/-5.2 |

\*Only perclinally dividing cells in the developing ligule were chosen for analysis

Periclinal divisions occur rarely in the maize epidermis (Poethig, 1984; Sharman, 1942), although they accounted for ~7% of in silico predicted divisions. We therefore examined cells from the developing ligule, an epidermally derived structure found in grass leaves, which contains a high proportion of periclinally dividing cells during specific developmental stages (Becraft et al., 1990; Sylvester et al., 1990). Micrographs of cells undergoing periclinal divisions within the developing ligule were selected for soap-film minimization predictions using Surface Evolver (n=34 cells, Figure 4A). While transverse divisions represented the global minimum division and also the most predicted division plane in leaf epidermal cells, the predicted periclinal divisions represented the global minimum in 97% of the cells in the developing ligule (n=33/34). Cells in this tissue expanded in the z-plane before the periclinal division was initiated (Figure 4B-C) suggesting that directional cell expansion rather than division plane specification may the driving force behind the eventual periclinal division in the developing ligule.

**FIG. 4.**
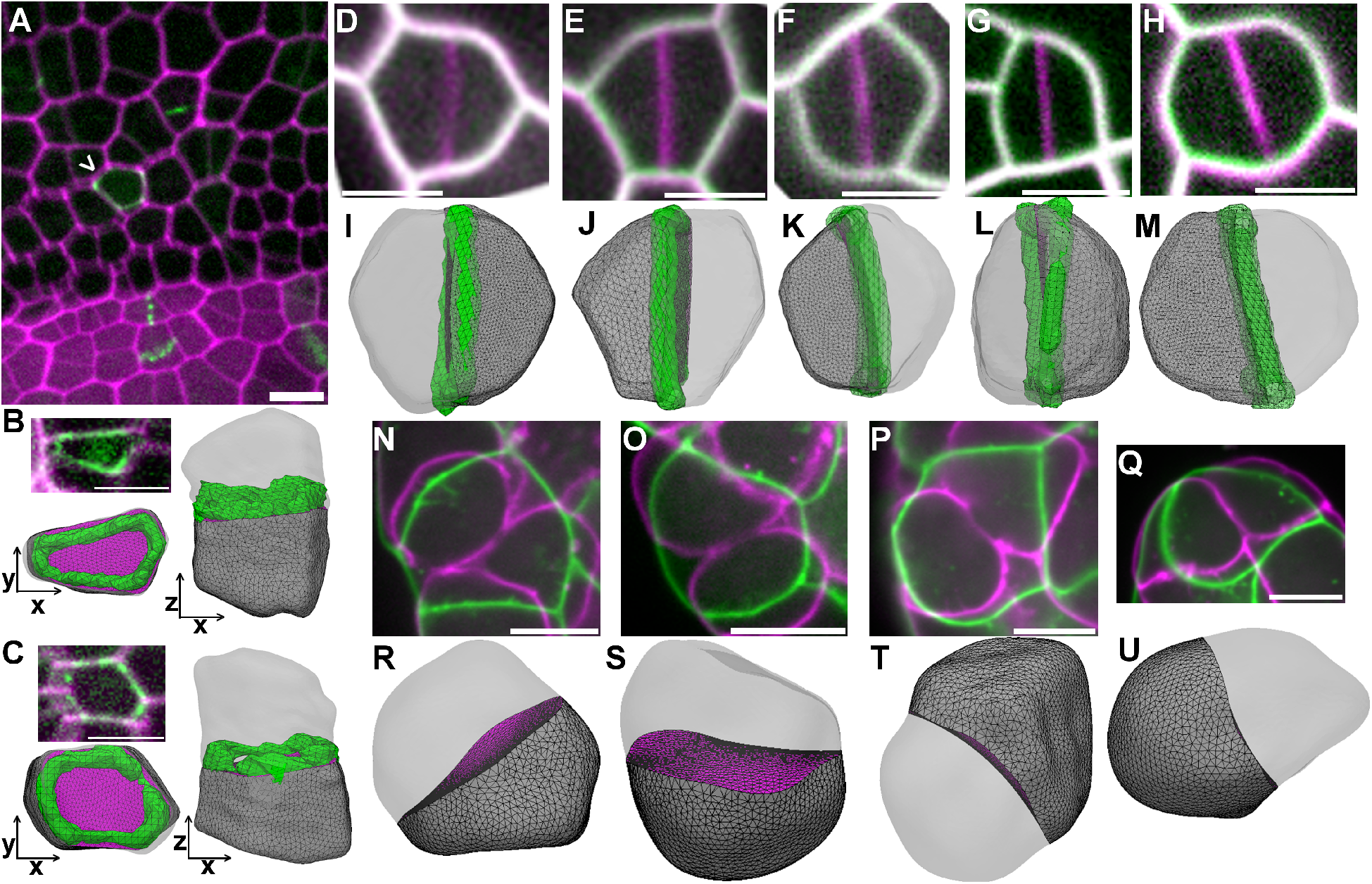
Division plane prediction in plant and animal cells. (A) Micrograph of maize developing ligule. Cell walls are stained with PI (magenta) and division sites are marked by TAN1-YFP (green). Arrowhead indicates periclinal division. (B-C) Micrographs of cells from the developing ligule expressing PIP2-CFP (magenta) for identification of cell outlines and TAN1-YFP (green) for division site location. The PPB and 3D reconstruction in both the XY (bottom left) and XZ plane (right) show the periclinal division plane. (D-H) Timelapse imaging of Arabidopsis guard cell division. The time point before the start of cytokinesis (green) overlaid with the completed division location (magenta). (I-M) 3D reconstruction of cells in (D-H) with the corresponding predicted division plane by Surface Evolver. The final division site was the newly formed cell wall (green) for comparison to the predicted division. (N-Q) Predicted division planes of cells in early gastrulation-stage *C. elegans* embryos. The cell shape prior to furrow ingression (green) was used for division predictions and overlaid with the dividing cell (magenta). (R-U) 3D reconstruction of cells in (N-Q) along with the corresponding predicted division plane by Surface Evolver. Scale bars are 10μm.

To test the generality of this method of division plane prediction, soap-film minima were predicted for *A. thaliana* symmetric guard mother cells and *Caenorhabditis elegans* embryonic cells. Time-lapse imaging of dividing guard mother cells in the *A. thaliana* epidermis was used to identify local minimum surface division planes. The final division plane was used as a marker for the division site to calculate offset between the predicted and in vivo divisions. Guard mother cell divisions that generated a guard cell pair were accurately represented by a soap film minimum predictions, with an average offset of 1.04 μm^2^ +/− 0.95 S.D. (Figure 4D-M), similar to offset observed in maize epidermal cells. Interestingly, the predicted division that matched the future in vivo location typically did not occupy the global minimum (n=7/8) but instead trended towards an anticlinal division plane with a larger surface area (Figure 4D-M). The cell that did not match this trend only had one anticlinal division class predicted, which was closely matched by the in vivo division. The eigenvalue ratio of these cells on average was 1.5 +/− 0.3 S.D., similar to (or even lower than) longitudinally dividing maize epidermal cells. Whether developmentally regulated cell shape parameters or mechanical cues promote long anticlinal divisions instead of the global minimum division is still unknown, although there are correlations between PPB location and local thickening of cell walls during guard cell division (Zhao and Sack, 1999). The close correspondence between a predicted division plane and the in vivo division indicates that soap film minimization accurately predicts the possible placement of divisions in flat, cylindrically-shaped cells in addition to rectangular prism-shaped cells.

Finally, *C. elegans* developing embryo cells were used to determine whether soap-film minima can be used to predict future division planes in symmetrically dividing animal cells. Time-lapse imaging was performed on dividing cells, and the cell shape immediately before furrow ingression was used for soap film minimization. While we could not accurately measure division plane offset in these cells due to their dynamic shape changes, the surface for one of the predicted division classes was approximately in the same location as the final division plane for the *C. elegans* cells (Figure 4N-U). The division class that matched the location of the in vivo division accounted for 75.8% +/− 28.4%; mean +/− S.D. of the surfaces generated by gradient descent surface minimization. These cells divided along the shortest division plane: the predicted division which matched the *in vivo* division was typically the global minimum for the cell (n = 7/8). These data provide evidence that the geometric cues contributed by pre-mitotic cell shapes can be used to predict the final division plane.

## Discussion

Much progress has been made recently on geometry-based modeling in 2D (Shapiro et al., 2015; Yoshida et al., 2014; Minc and Piel, 2012; Besson and Dumais, 2011; Dupuy et al., 2010; Sahlin and Jönsson, 2010) but the transition to 3D plant cell modeling is essential to understanding the fundamental mechanisms of division plane orientation (Chakrabortty et al., 2018a; Yoshida et al., 2014). Symmetric division planes can be predicted by importing any 3D cell shape into Surface Evolver (Brakke, 1992) and performing iterative gradient descent from multiple starting planes to generate local soap-film minima. We demonstrated that this model generates reasonable predictions for three different maize cell types, as well as *A. thaliana* and *C. elegans* cell divisions. Modeling each cell produced one or more local minima, that when compared to in vivo division, typically produced a close match.

In addition to predicting multiple potential division plane orientations, the predicted divisions were directly compared to the location of the PPB, a structure known to predict the future division site (Smertenko et al., 2017; Rasmussen et al., 2013; Rasmussen and Bellinger, 2018). Several elegant studies used time-lapse imaging to compare cells before and after division (von Wangenheim et al., 2016; Shapiro et al., 2015; Louveaux et al., 2016) but they did not directly assess PPB location in comparison to predicted divisions. This step allowed us to identify alterations in PPB location compared to the mathematically optimal division. When PPB locations shifted, it was due to coordination between cells to prevent the formation of a four-way-junction. Four-way junction avoidance is a well-known feature of plant cells (Flanders et al., 1990; Gunning et al., 1978; Sinnott and Bloch, 1941), but it was not known whether PPB localization was driven by cell shape or interactions between cells. Although previously suspected, we demonstrated that the PPB is offset from the soap-film predicted division site during four-way junction avoidance. Alterations in PPB location to avoid an adjacent wall or another PPB, strongly suggests that a signal, whether chemical, mechanical or both, is actively communicated between cells. Therefore, PPB dynamics and positioning are likely modulated by cell-cell interactions.

It is not surprising that the overall probabilities predicted by our model, which are generated via unbiased soap film minimization by sampling the entire cell shape, do not describe in vivo proportions of division planes observed in maize epidermal cells, particularly the underprediction of longitudinal divisions and overprediction of periclinal divisions. During symmetric cell division, cells can divide with an increased probability along a longer plane rather than the shortest plane, such as when they are under mechanical stress (Lintilhac and Vesecky, 1984; Louveaux et al., 2016; Landrein and Hamant, 2013). In addition, the division axis of asymmetrically dividing cells can be influenced by the growth axis of the tissue and accompanying mechanical stresses (Bringmann and Bergmann, 2017). The cortical microtubule array interacts with and helps direct formation of the cell wall polymers such as cellulose, leading long-term to alterations in cell shape (Wasteneys and Ambrose, 2009; Paredez et al., 2008; Zhu et al., 2015; Li et al., 2012). Cortical microtubule arrays and sometimes subsequent division planes align parallel to plane of stress (Landrein and Hamant, 2013; Lynch and Lintilhac, 1997; Lintilhac and Vesecky, 1984; Asada, 2013) and after wounding or neighbor cell ablation (Heisler et al., 2010; Hush and Overall, 1996; Lintilhac and Vesecky, 1981; Sinnott and Bloch, 1941). In the vast majority of divisions we observed, cell shape predicts all potential division planes, but local and tissue level stresses likely alter the final location, similar to microtubule array realignment after mechanical perturbation (Sampathkumar et al. 2014).

Epidermal cells divide almost exclusively along an anticlinal plane (Poethig, 1984; Galletti et al., 2016), even though soap-film predictions indicate that periclinal divisions are possible, suggesting that molecular or mechanical mechanisms inhibit periclinal divisions in the epidermis. Epidermal cell fate is specified early, and multiple gene regulatory networks uniquely define and specify the epidermal cell layer (Takada and Iida, 2014). In addition, the outer epidermal wall is both covered in cuticle and typically much thicker than interior walls in maize (Carpita et al., 2001) and tomato (Kierzkowski et al., 2012). The epidermis is thought of as a load-bearing layer under tension: its growth influences the overall size of the tissue (Marcotrigiano, 2010) and it acts as a shell resisting MPa turgor pressure (Beauzamy et al., 2015). Although epidermal cells do not typically divide along a periclinal plane, the mechanisms that normally prevent periclinal divisions are still unknown.

Periclinal divisions tend to occur during developmental processes such as formation of the vasculature, root, anther and ligule (Walbot and Egger, 2016; Sylvester et al., 1990; Wachsman et al., 2015; Van Norman, 2016; Sakaguchi and Fukuda, 2008; Poethig, 1984; Lang Selker and Green, 1984). Therefore, we chose to analyze cells undergoing periclinal divisions during ligule development in maize. During ligule development, the periclinal division is the global minimum prediction, suggesting that these cells expanded before the division plane was established. The potential coordination between expansion and division plane orientation was revealed by the Arabidopsis BREAKING ASYMMETRY IN THE STOMATAL LINEAGE (BASL) protein, which localizes to one edge of the cell to promote an asymmetric division during stomatal development. Interestingly, ectopic BASL expression promotes localized and directional expansion, dependent on Rho of Plants (ROP) monomeric GTPase function (Dong et al., 2009). Directional cell expansion may be a general mechanism to promote division plane orientation along a specific plane.

Our model has the potential to identify the correct placement of the symmetric division plane in cells with any shape. Defects in cortical microtubule organization or other defects can lead to aberrantly shaped daughter cells (Komis et al., 2017; Kirik et al., 2012; Hashimoto, 2015; Pietra et al., 2013; Martinez et al., 2017). In mutants with aberrant cell shapes, it can be difficult to determine whether the symmetric division plane specified by the PPB follows the geometry of the cell (Lipka et al., 2014; Martinez et al., 2017; Pietra et al., 2013; Mir et al., 2018). Potentially “misplaced” PPBs may arise either as a direct consequence of altered cell shape, the absence of protein function, or indirectly through alterations in stress or tension on the cell (Willis et al., 2016). This model may be useful in uncoupling cell shape defects from division plane specification defects.

Cell shape changes during animal cell mitosis make it difficult to assess whether factors such as geometric cues before mitosis are used to orient the division or if the division plane is specified by other interactions as the cell is dividing (Minc and Piel, 2012). Surface area minimization (akin to soap bubbles) along with cell-cell contacts or adhesion have been used to describe patterning and division plane orientation of animal cells (Kafer et al., 2007; Hayashi and Carthew, 2004; Goldstein, 1995; Pierre et al., 2016; Gibson et al., 2011). Landmark cues including tricellular junctions during mitotic rounding and contacts with the extracellular matrix can be maintained to properly orient animal cell division planes (Bosveld et al., 2016; Théry et al., 2005). While geometric cues are only one of several factors promoting division plane orientation in plant and animal cells, soap-film surface area minimization accurately predicts in vivo symmetric division planes regardless of specific, and as yet still mostly unknown, mechanisms.

## Materials and Methods

### Imaging of Maize Tissue

Maize leaves from a 28 day-old plant expressing a live cell marker for microtubules were dissected to reveal symmetrically dividing cells (Sylvester, 2000). YFP-TUBULIN (YFP-variant Citrine fused to α-TUBULIN (GRMZM2G153292), (Mohanty et al., 2009) was used to identify PPB location. Cell walls were stained with 0.1 mM propidium iodide (Fisher) or plasma membranes were identified using PIP2-1-CFP (PLASMA MEMBRANE INTRINSIC PROTEIN2-1 (GRMZM2G014914) fused to Cerulean Fluorescent protein (Mohanty et al., 2009). Two distinct leaf developmental stages were chosen for analysis, a young leaf, L14, and an older leaf, L9 or L10. Leaf 14 (total blade height = 15.6mm) from a 28 day-old plant was dissected and the abaxial side was imaged near the leaf margin. Leaf 9 or 10 (ligule height ~2 mm) was imaged directly above the ligule near the margin. They were loaded into a Rose chamber for imaging on an inverted Nikon Ti stand with Yokogawa spinning disk and a motorized stage (ASI Piezo) run with Micromanager software (micromanager.org) and built by Solamere Technology. Solid-state lasers (Obis) and emission filters (Chroma Technology) used excitation, 445; emission, 480/40 (for CFP-TUBULIN and PIP2-1-CFP); excitation, 561; emission, 620/60 (for propidium iodide); and excitation, 514; emission, 540/30 (for YFP-TUBULIN and TAN1-YFP). Perfluorocarbon immersion liquid (RIAAA-678; Cargille) was used for 40× or 60× water-immersion objectives with 1.15 and 1.2 numerical aperture, respectively. Micrographs of transverse or longitudinal PPBs and cell wall or membranes in 0.2 or 0.4 μm Z-stack increments. The Z-stack images were opened in FIJI (http://fiii.sc/) and then cropped to one cell. The cell wall or plasma membrane outline was extracted using the following process in FIJI: segmentation, find maxima and make binary (Schindelin et al., 2012). The PPB was extracted using trainable Weka segmentation in FIJI (Arganda-Carreras et al., 2017). After manual correction of cell wall outlines and PPB binary files, the cell wall and PPBs were converted into a 3D surface using *Plugin ≫ 3D Viewer* with a resampling factor of 2. Then, the 3D surface was exported by using the options *File ≫ Export As ≫ STL (ASCII)*. Maize leaves were dissected to reveal preligule bands where periclinal divisions were frequent (L10 PLB height ~ 2mm). Propidium iodide, PIP2-CFP, YFP-TUBULIN, and TAN1-YFP were used identify cell outlines and division site locations.

### Time-lapse imaging of maize tissue

The adaxial side of maize leaf blade near the margin was imaged at 21 °C. For time-lapse experiments, leaf samples were mounted on a coverslip within a Rose chamber, surrounded by vacuum grease,then covered with another coverslip inside the chamber (<u>Rasmussen 2016</u>). Maize epidermal cells expressing YFP-TUBULIN were imaged at 1 μm Z intervals every 5 minutes for up to 6 hours, and viability was assessed by presence of actively dividing cells. Cell outlines and PPBs were extracted using the YFP-TUBULIN signal and segmented using Weka Trainable Segmentation algorithm (Arganda-Carreras et al., 2017). Timelapse imaging was also conducted using maize expressing TAN1-YFP and CFP-TUBULIN (Cyan fluorescent protein fused to β-TUBULIN, GRMZM2G164696) to identify mitotic stages.

### Imaging of Arabidopsis Guard cell divisions

Confocal imaging of 4-day-old light-grown Arabidopsis thaliana seedlings of the Col-2 ecotype, expressing LTI6b-GFP (Cutler et al., 2000 - PMID 10737809) was performed on a Zeiss Axio Observer microscope with a Yokogawa CSU-X1 spinning disk head and a 63X oil immersion objective, and a 488-nm excitation laser and a 525/50-nm emission filter. Z-stack time-lapse (stacklapse) imaging was performed as in (Peterson and Torii, 2012 - PMID 23299369): a cotyledon was cut from a seedling and placed under a 0.5% agar pad in a NuncTM Lab-TekTM coverglass chamber (Thermo Fisher; catalog no. 155360). The edge between the agar pad and the chamber was sealed by adding additional melted agar. The abaxial side of the cotyledon was oriented facing the coverglass. The chamber was mounted on the microscope stage with the lid on to minimize evaporation. Stacklapse images were collected with an interval of 0.5 h, up to 48 h, and a step size of 0.5 μm using 2% 488-nm laser power and 200-ms exposures.

### Imaging of *C.elegans* embryo cell divisions

*C. elegans* animals were cultured on Normal Growth Media plates, fed Escherichia coli (OP50 strain), and grown at 20°C. The strain used is LP733 *cpIs131* [*Pmex-5*>mScarlet-I-C1 ::PH::*tbb*-2 3’UTR lox N ttTi5605] II. Embryos were imaged using methods described in (<u>Heppert et al. 2018</u>). In brief, embryos were mounted at approximately the 24-cell stage and imaged on a spinning disk confocal microscope. mScarlet was excited using a 561-nm solid state laser with a 568LP emission filter set. Single channel embryo samples were filmed at 30-sec intervals. Images were cropped and rotated, and brightness and contrast were adjusted using Fiji.

### Sample Size

Two individual maize leaves from two separate plants were carefully imaged to capture developmental “snapshots” of all the cell divisions within those tissues. The purpose of this experiment was to capture enough cell shape and division plane data to compare these two different developmental stages. 67 cells were captured from a developmentally younger leaf (leaf L14, leaf height 16 mm from a 28 day old maize plant) and 114 cells were captured from a developmentally older leaf were used (L9 or L10, ligule height ~2 mm from a 28 day old maize plant). For time-lapse imaging data, maize leaf epidermal cells from three individual maize plants were analyzed. Cells from maize ligule cells, (n = 34) were captured from more than three individual ligules from three separate 28 day old maize plants (L10 pre-ligular band height ~ 2 mm). *Arabidopsis thaliana* guard cell divisions (n = 8) were captured from 6 individual cotyledons of different plants (4-day-old light-grown seedlings, Col-2 ecotype). *C. elegans* cell divisions (n = 8) were captured from two individual *C. elegans* embryos (at ~24-cell stage).

### Surface Evolver

The 3D surfaces were imported into Surface Evolver. Spherical harmonics, a common 3D rendering technique to generate a formula to describe complex cell shapes, were generated in Surface Evolver. Spherical harmonics approximate the cell outline as a function of spherical coordinate angles (Givoli, 2004). A Github workflow diagram for the Surface Evolver can be found at (https://github.com/idhayes/predictive_division/blob/master/doc/predictive_division.svg). Cells from maize leaf and ligule, and Arabidopsis guard mother cells were smoothed using 30th degree spherical harmonics. For modeled *Caenorhabditis elegans* embryo cells, 10th degree spherical harmonics were used to approximate the cell shapes. Spherical harmonics code in Surface Evolver https://github.com/jdhayes/predictive_division/blob/master/etc/SphericalHarmonics.cmd. Next, soap film minimization was performed using 241 initial planes (https://raw.githubusercontent.com/idhayes/predictive_division/master/etc/241points.txt) with normals uniformly distributed over a sphere (https://github.com/idhayes/predictive_division/blob/master/etc/trailer.inc). Gradient descent, a computational method to generate a soap-film minimum surface, generated surface areas reflecting predicted divisions. Predicted divisions with similar surface areas were grouped together and categorized by their location in the cell relative to the tissue as transverse, longitudinal, periclinal and other. To determine the PPB offset for each cell, the midplane of the PPB was aligned to the predicted division. The surface with the lowest offset from the in-vivo location of the PPB was used in Figure 3 (code at https://github.com/idhayes/predictive_division/blob/master/etc/PPB_OFFSET.txt). This measurement indicates how closely the PPB and the predicted division align. Cell shape was described using eigenvalue ratios. Eigenvalue ratios were calculated using the following steps. First, using Surface Evolver, three eigenvectors were calculated from the moment of inertia matrix for each cell. Second, the square root was taken of the largest eigenvector divided by the smallest eigenvector to generate the final eigenvalue ratio. Second, the square root was taken of the largest eigenvector divided by the smallest eigenvector to generate the final eigenvalue ratio (https://raw.githubusercontent.com/idhayes/predictive_division/master/etc/inertia.txt).

### Spherical Harmonics

The 3D surfaces generated from FIJI were imported into Surface Evolver. Since Surface Evolver input files need to be represented by a formula, surfaces were defined using 30th degree spherical harmonics. Increasing spherical harmonic degrees increased cell shape accuracy, but also increased computational time. 30th degree spherical harmonics provided accurate cell shape while minimizing computational time. Using 10th degree spherical harmonics led to inaccurate cell shapes which lacked many of the distinguishing geometric features of cells (Fig. S1A-E) resulting in overprediction of transverse divisions for the cells (Table S1). 20th degree spherical harmonics did not properly model the corners and some features of the cell shape (Fig. S1A-E). 30th degree spherical harmonics more accurately represented the original cell shape, and allowed for efficient and complete calculation of gradient descent (Fig. S1A-E). Results for predicted divisions of different spherical harmonic degrees for each cell are summarized in Table S1.

## Acknowledgments

CH was supported by the National Science Foundation (NSF) Research Experience for Undergraduates grant #1461297 to UC Riverside Center for Plant Cell Biology (CEBCEB). CGR gratefully acknowledges NSF-MCB #1505848 and #1716972 and a collaborative seed grant to CGR and AM from the UCR Office of Research. Work by CTA and YR was supported by NSF-MCB #1616316.

---

**Figure S1.**
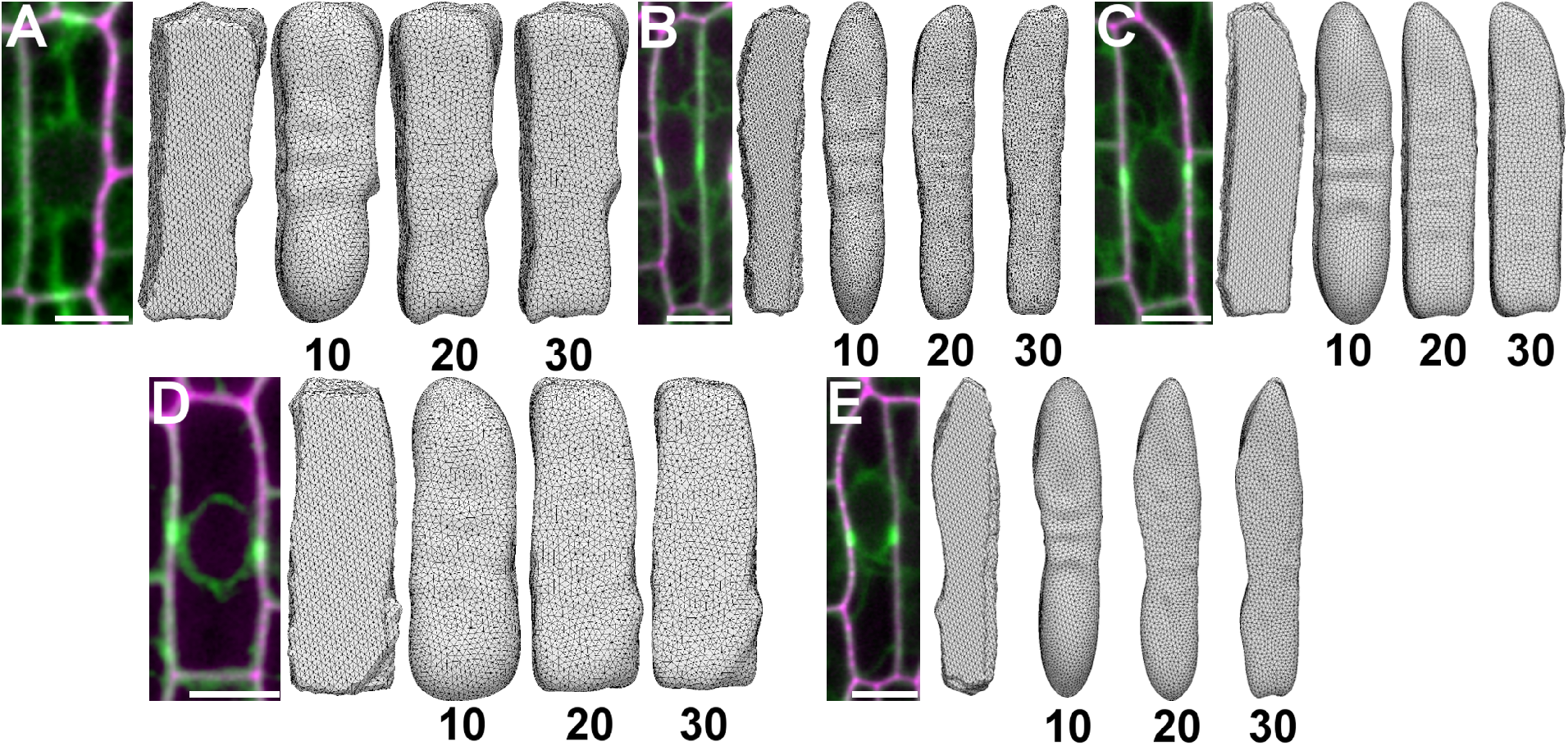
Spherical Harmonics: different degrees. (A-E) Micrograph of maize epidermal cells expressing YFP-TUBULIN (green) and stained with propidium iodide (magenta). Left-most 3D representation is the original cell threshold without spherical harmonics applied. Proceeding from left to right 10th, 20th and 30th degree spherical harmonics (labeled below). Scale bar is 10μm.

**Table S1.** Analysis of Cells at different Degrees of Spherical Harmonics

| Cell | Degree of SH | Transverse (%) | Longitudinal (%) | Periclinal (%) | Other (%) |
| --- | --- | --- | --- | --- | --- |
| **Figure S1A** | 10 | 92.1 | 0 | 7.9 | 0 |
|  | 20 | 88.8 | 2.5 | 8.7 | 0 |
|  | 30 | 86.7 | 3.7 | 9.6 | 0 |
| **Figure S1B** | 10 | 100 | 0 | 0 | 0 |
|  | 20 | 90 | 0 | 10 | 0 |
|  | 30 | 91.3 | 0 | 8.6 | 0 |
| **Figure S1C** | 10 | 100 | 0 | 0 | 0 |
|  | 20 | 92 | 0 | 8 | 0 |
|  | 30 | 91.2 | 0 | 8.7 | 0 |
| **Figure S1D** | 10 | 100 | 0 | 0 | 0 |
|  | 20 | 91.3 | 5 | 3.7 | 0 |
|  | 30 | 90 | 4.2 | 5.8 | 0 |
| **Figure S1E** | 10 | 100 | 0 | 0 | 0 |
|  | 20 | 94.6 | 0 | 5.4 | 0 |
|  | 30 | 92 | 0 | 8 | 0 |

